# Left parietal tACS at alpha frequency induces a shift of visuospatial attention

**DOI:** 10.1101/644237

**Authors:** T. Schuhmann, S. K. Kemmerer, F. Duecker, T.A. de Graaf, S. ten Oever, P. De Weerd, A.T. Sack

## Abstract

**Background:** Voluntary shifts of visuospatial attention are associated with a lateralization of occipitoparietal alpha power (7-13Hz), i.e. higher power in the hemisphere ipsilateral and lower power contralateral to the locus of attention. Recent noninvasive neuromodulation studies demonstrated that alpha power can be experimentally increased using transcranial alternating current stimulation (tACS).

**Objective/Hypothesis:** We hypothesized that tACS at alpha frequency over the left parietal cortex induces shifts of attention to the left hemifield. However, spatial attention shifts not only occur voluntarily (endogenous), but also stimulus-driven (exogenous). In order to study the task-specificity of the potential effects of tACS on attentional processes, we administered three conceptually different spatial attention tasks.

**Methods:** 36 healthy volunteers were recruited from an academic environment. In two seperate sessions, we applied either high-density tACS at 10Hz, or sham tACS, for 35-40 minutes to their left parietal cortex. We systematically compared performance on endogenous attention, exogenous attention, and stimulus detection tasks.

**Results:** In the Endogenous attention task, we found a greater leftward bias in reaction times during left parietal 10Hz tACS as compared to sham. There were no stimulation effects in the exogenous attention or stimulus detection task.

**Conclusion:** The study shows that high-density tACS at 10Hz can be used to modulate visuospatial attention performance. The tACS effect is task-specific, indicating that not all forms of attention are equally susceptible to the stimulation.

## Introduction

Visual scenes typically include many stimuli. As our brains are not able to efficiently process all stimuli simultaneously, some need to be prioritized over others. Such selection can be achieved with visuospatial attention, which enables preferential processing of stimuli at a location of interest, as shown by decreased reaction times (RT) [1]. To some extent, we naturally display preferential processing in one hemifield relative to the other [2]; we have a spatial attention *bias*. On top of this, spatial attention can be shifted, either voluntarily (endogenous), or automatically, when captured by a salient stimulus (exogenous). Different tasks have been used to assess these processes. *Visual detection tasks* measure low-level perceptual sensitivity and attentional selection, also in the context of multiple, simultaneously presented stimuli. They can reveal information about attentional biases, but do not include an attentional manipulation and therefore leave higher-order attentional processes out of consideration. In contrast, *orienting tasks* (endogenous and exogenous) directly capture higher-order attentional processes, by comparing the efficiency of attention shifts across various cue conditions [3]. In endogenous attention shifts, participants voluntarily allocate attention based on internal goals or task instructions. Exogenous attention shifts on the other hand are stimulus-driven and automatic. Electroencephalography (EEG) studies show that a shift in endogenous attention is associated with an occipitoparietal alpha (8-12Hz) power lateralization, i.e. alpha power increases in the ipsilateral relative to the contralateral side of attention [4–7], Even on a trial-by-trial basis, alpha power lateralization predicts RT to stimuli in either hemifield [7]. Interestingly, while endogenous attention shifts are associated with a lateralization of occipitoparietal alpha power, evidence for a similar link between exogenous attention and alpha power lateralization is still lacking.

Hemineglect patients show a pathological attention bias, marked by reduced responses to stimuli in the contralesional hemifield, generally caused by (usually right-hemispheric) unilateral stroke in the temporoparietal lobe [8,9]. Patients are slower and less accurate in contralesional target detection [10,11], and display a spatial orienting bias in endogenous and exogenous tasks [12–14], In EEG, the amplitude [15–17] and amplitude variability [15] of alpha oscillations are reduced over their damaged hemispheres. In the recovery period, alpha power increases again [18] and is associated with clinical improvement [17,19,20]. These results suggest that alpha power is related to attentional bias and orienting performance.

To empirically demonstrate the causal relevance of parietal alpha oscillations in attention, one should modulate alpha power experimentally. This can be achieved with non-invasive brain stimulation (NIBS) techniques such as transcranial alternating current stimulation (tACS). TACS consists of low-intensity electrical current flowing rhythmically back and forth between two (or more) electrodes [21–23], Recent studies combining tACS with EEG show that tACS at alpha frequency leads to an elevation of alpha power [22–25]. TACS has been used to studv the functional role of oscillations in various coqnitive processes in healthv volunteers [26–31]. Unilateral tACS at alpha frequency has previously been shown to influence spatial attention performance [30,31]. Left temporocentral alpha tACS induced a leftward bias in an auditory attention/working memory task [30]. In the visual domain, right parietal alpha tACS modulated visuospatial attention performance in a landmark task (experiment 1, [31], although this finding could not be replicated (experiment 2, [31]).

In the current study, we investigated whether left parietal tACS at alpha frequency results in a significant shift of visuospatial attention to the left hemifield in healthy volunteers. We used a high-density ring electrode montage over the left parietal cortex targeting the left hemispheric attention network. Considering the conceptual difference between simple detection versus orienting tasks and endogenous versus exogenous spatial attention shifts, we evaluated whether alpha tACS affected these processes differentially. We stimulated the left parietal cortex at alpha frequency (10Hz) and sham in separate sessions and measured visuospatial attention performance in a visual detection, endogenous attention, and exogenous orienting spatial attention task. We hypothesized that left parietal 10Hz tACS induces an attentional leftward bias relative to sham in the endogenous attention and detection task. As it is still unknown whether parietal alpha oscillations are also associated with exogenous attention shifts we had no a priori hypothesis regarding the effect of tACS on the Exogenous attention task.

## Methods

### Participants

We tested 36 healthy, right-handed students with normal or corrected to normal vision (18 women, mean age = 21.56 years, age range = 18-29 years). At the beginning of each session, participants gave their written informed consent and were screened for tACS safety. For this, we followed the recommended procedures of Antal and colleagues [32], screening for e.g. skin diseases, neurological disorders, implants, pregnancy and medication.

### Procedure

Each participant underwent 10Hz as well as sham tACS in two separate sessions. A session started with practicing the detection, endogenous attention and exogenous attention task. TACS was subsequently applied at either 10Hz or sham during which participants performed the visuospatial attention tasks. In each session, participants performed all three visuospatial attention tasks in a counterbalanced order. TACS never exceeded 40 minutes and was switched off after completion of all tasks. An eye tracker was used for the Endogenous and Exogenous attention task to record eye movements.

### Blinding

Throughout the experiment, participants were blinded to the experimental hypotheses and the stimulation protocol. At the end of each session, we administered a questionnaire which required the participants to judge whether real or sham stimulation was applied. To assure that participants were not able to differentiate between the two stimulation conditions, we ran a generalized estimating equation analysis [33] with actual stimulation condition (real or sham) as factor and rated stimulation condition as dependent variable (table 1). Rated stimulation condition was assessed on an ordinal scale with seven levels. The value one corresponded to ‘I definitely experienced placebo/sliam stimulation’ and the value seven to ‘I definitely experienced real stimulation’. According to the Wald chi square test, the actual stimulation condition did not affect the rated stimulation condition (*X*^2^(1, N=64)=.205, p=.651), indicating that blinding was indeed maintained.

**Table 1.**
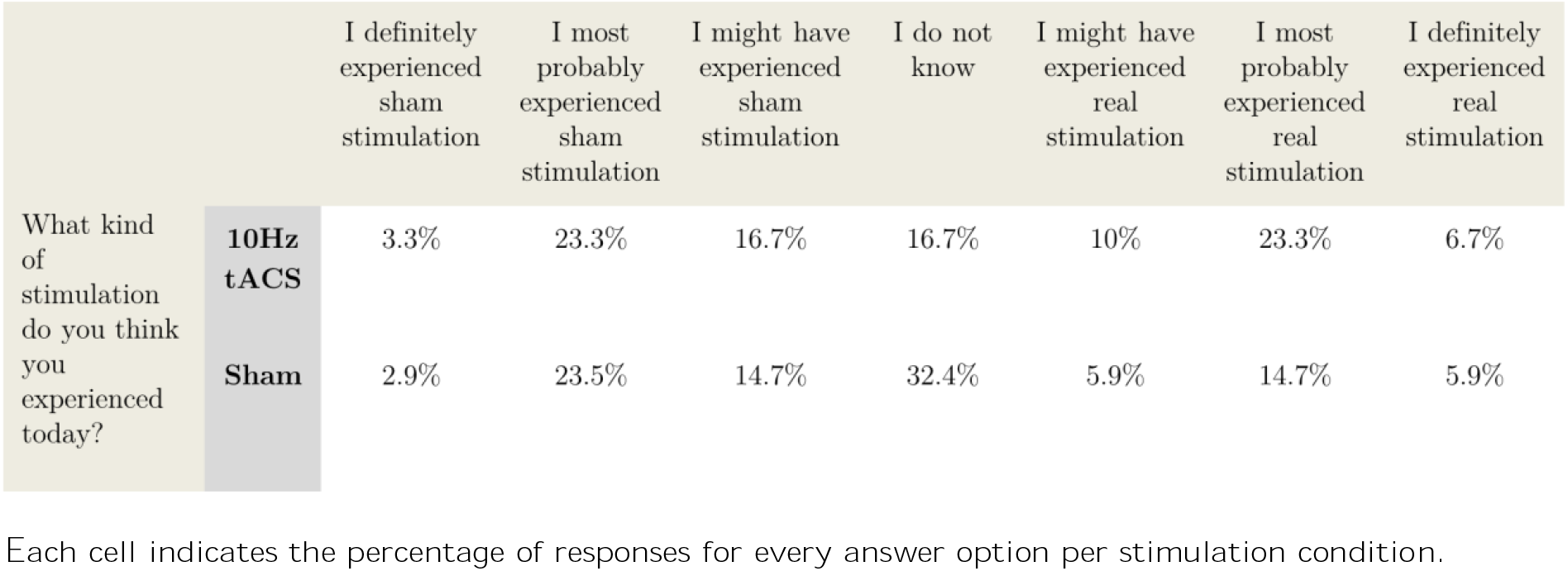
Outcomes of a post-stimulation questionnaire assessing the rated stimulation condition based on the participant’s subjective experience.

|  |  | I definitely experienced sham stimulation | I most probably experienced sham stimulation | I might have experienced sham stimulation | I do not know | I might have experienced real stimulation | I most probably experienced real stimulation | I definitely experienced real stimulation |
| --- | --- | --- | --- | --- | --- | --- | --- | --- |
| What kind of stimulation do you think you experienced today? | **10Hz tACS** | 3.3% | 23.3% | 16.7% | 16.7% | 10% | 23.3% | 6.7% |
|  | **Sham** | 2.9% | 23.5% | 14.7% | 32.4% | 5.9% | 14.7% | 5.9% |

Each cell indicates the percentage of responses for every answer option per stimulation condition.

### TACS

A small circular (diameter: 2.1cm, thickness: 2mm) and a large (Outer diameter: 11cm; Inner diameter: 9cm, thickness: 2mm) rubber ring tACS electrode (NeuroConn, Ilmenau, Germany) were placed onto the left parietal cortex, with the small electrode positioned over P3 and the large electrode centred around it (figure 1A). This ring electrode montage enables a higher spatial focality as compared to standard rectangular electrodes [34]. Conductive gel (en20 paste, Weaver and Company, Aurora, CO, USA) was applied between skin and electrodes to reduce the impedance to below lOkΩ. Stimulation frequency and intensity were set to 10Hz and 1mA peak to peak, phase offset was set to 0 and 100 cycles were used for ramping up. The sham stimulation was ramped up and then immediately ramped down with each 100 cycles.

**Fig 1.**
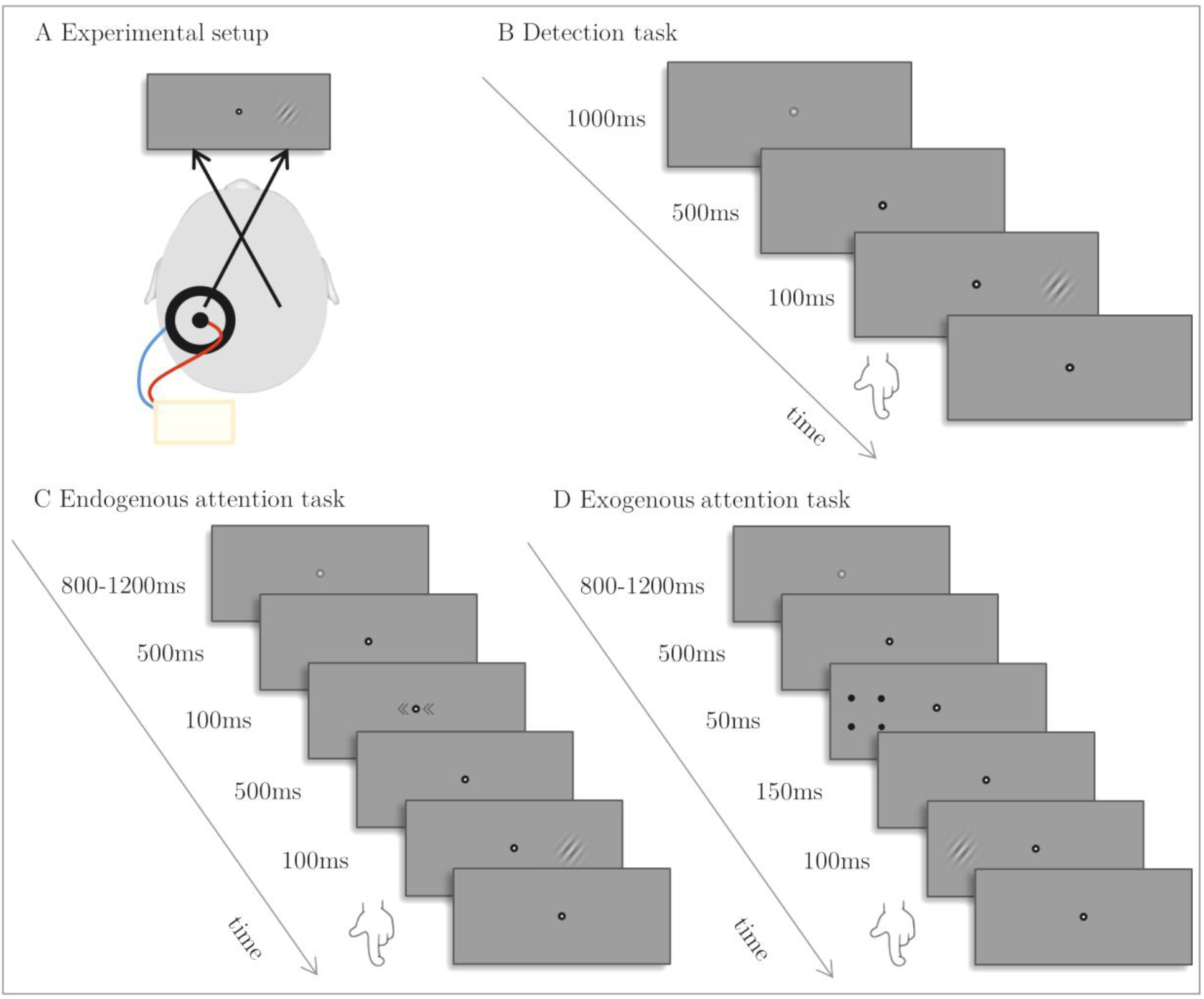
*Experimental setup and example trials from the three attention tasks. A TACS setup*. TACS ring electrodes were centered on P3 and each participant was stimulated at 10Hz or sham in separate sessions. B *Example trial from the Detection task*. A sinusoidal grating was presented either in the left, right or both hemifields. Participants had to indicate the location of the sinusoidal grating. C *Example trial from the Endogenous attention task*. A trial started with the presentation of a fixation point followed by an endogenous cue (<<·<<, >>·>> or <<·>>) directing attention to the left, right or both sides. Thereafter, a sinusoidal grating tilted 45° to either side was shown in the left or right hemifield. The participants had to indicate whether the grating was turned to the left or right by pressing the corresponding button (invalid trial in this example). D *Example trial from the Exogenous attention task*. Similarly to the Endogenous attention task, a trail started with the presentation of a fixation point followed by an exogenous cue. The exogenous cue consisted of four black dots forming a square and surrounding either the left or right potential target location (directional cue) or a luminance change of the background color of the screen (neutral cue). Then the sinusoidal grating was presented in the left or right hemifield and participants had to discriminate its orientation.

### Visuospatial attention tasks

The detection task was preceded by a short calibration procedure during which the contrast of a sinusoidal grating was manually reduced to slightly above the participant’s detection threshold. This value was used as an initial contrast value for the detection task. During the detection task, sinusoidal gratings with randomized left or right orientation were presented in the left, right, or both hemifields (figure 1B). The participant had to indicate the location of the stimulus and the contrast of the stimuli in the left, right or both hemifields were adapted according to a staircase algorithm aiming at 50% accuracy [35]. One iteration of the detection task comprised three staircases of each 40 trials resulting in three contrast thresholds per session (one for left, one for right and one for the bilateral targets). In total, the detection task took approximately 10 minutes.

The Endogenous attention task started with the presentation of a fixation point, which changed in greyscale after a jittered interval. Then two arrowheads pointing to the left (<<·<<), right (>>·>>) or both sides (<<·>>) flanking the fixation point and predicting the correct target location with 80% validity were shown. This was followed by a sinusoidal grating with a Gaussian envelope, which appeared at 7° eccentricity either in the left or right hemifield (figure 1C). The grating was rotated by 45° in either clockwise or counterclockwise direction and the task of the participant was to discriminate its orientation. There were in total 336 trials consisting of 192 valid trials, 48 invalid trials, and 96 neutral trials and task duration was approximately 20 minutes.

The same fixation point and target stimulus were also used for the Exogenous attention task (figure 1D). As opposed to the endogenous version, the cues consisted of a change in background luminance (neutral cue) or four dots surrounding one of the possible target locations and indicating the correct target location with 50% validity (directional cue). The sinusoidal grating was presented at 14° eccentricity and the participant had to discriminate its orientation. The Exogenous attention task consisted of 216 trials in total with 72 trials per type of cue (valid, neutral, invalid), lasting approximately 10 minutes in total.

Each cell indicates the percentage of responses for every answer option per stimulation condition.

A more detailed description of the tasks can be found in Duecker and colleagues [3]. All tasks were presented with 60Hz on a gamma-corrected liyama ProLite monitor at 57-cm viewing distance. Video mode was 1920×1080 and the background luminance was 100cd/m^2^. The software application presentation (NeuroBehavioural Systems, Albany, CA) was used for running the tasks interfacing with MATLAB for the staircase algorithm [35].

### Eye tracker

Eye tracking (Eyelink1000, SR Research, Mississauga, Ontario, Canada) was used during the endogenous and exogenous cueing task. Initially a 9-point calibration and validation procedure was executed. Then we used monocular eye tracking at 1000Hz to track gaze position sample by sample point. Participants were told to keep their chin in the chin rest at all times to avoid movements. Post-hoc trials containing eye blinks and eye movements exceeding 2° of visual angle in the time window from 100ms before cue onset until stimulus onset were deleted (5.92% of all trials for the endogenous and 2.65% for the exogenous version). No eye tracking was used for the detection task, as it does not include a (re)orienting component.

### Statistical analysis

For both the Endogenous and Exogenous attention tasks we removed trials with extreme RTs based on the median +/− 1.5*interquartile range (IQR) criterion. To assure a sufficient number of trials per cell we calculated the average amount of trials over both hemifields per *Stimulation condition* and *Type of cue*. Only participants with more than 15 trials per stimulation condition and cue type were included in the analysis.

We performed repeated-measures analyses of variance (RM-ANOVA) to compare the condition averages of median RTs. A RM-ANOVA on sham tACS data, with factor Cue-Type, validated our attention tasks. For the detection task, a RM-ANOVA with factors *Stimulation condition* and *Stimulus location* compared contrast thresholds. To test extinction-like effects in the 10Hz tACS condition in incorrect bilateral trials, we performed a RM-ANOVA with *Stimulation condition* and *Indicated location of the stimulus* (left or right) as factors and number of trials as dependent variable.

For the Exogenous attention task, median RTs based on only correct trials were computed per *Hemifield*, *Type of cue* and *Stimulation condition*. Then, the visuospatial attention bias was calculated by subtracting the RTs to right-hemifield stimuli from RTs to left-hemifield stimuli. Resulting attention bias scores were compared in RM-ANOVA with factors *Type of cue* and *Stimulation condition*. The same main analysis was done for the Endogenous attention task, followed by two post-hoc analyses. We collapsed the median RTs across the three levels of cue and ran a Repeated Measures ANOVA with *Hemifield* and *Stimulation condition* as factors. With a t-test, we also tested whether there is the difference between RTs in the left and right hemifield in the 10Hz stimulation condition relative to sham.

## Results

Average accuracy over both hemifields was 93% (range: 69%-100%) for the Endogenous attention task. Two participants were not included in this analysis because of an insufficient number of correct trials without eye artefacts. For the Exogenous attention task, average accuracy over both hemifields was 93% (range: 67%-100%). Because of an insufficient number of correct trials without eye artefacts, one participant was excluded from the analysis. All participants were included in the analysis of the visual detection task.

### Cueing effects in the Endogenous and Exogenous attention task

First, we analyzed the data of the Endogenous attention task as acquired in the sham session and tested whether the manipulation of attention with the endogenous cues was successful. We ran a Repeated Measures ANOVA with median RT averaged over both hemifields as dependent variable and *Type of Cue* as factor. Establishing the cueing effect (figure 2A) we found a main effect of *Type of Cue* (F(2,66)=35.33, p<0.001), with significantly slower RTs in invalid trials (M=534.15, SEM=10.02) as compared to neutral (M=513.32, SEM=9.25) (t(33)=4.23, p<.001) and valid trials (M=493.35, SEM=7.95) (t(33)=7.04, p<.001) and significantly faster RTs in valid as compared to neutral trials (t(33)=−5.58, p<.001) (figure 2A).

The same analysis was done for the Exogenous attention task. Also here, there was a main effect of *Type of cue* (F(2,68)=29.58, p<.001) with faster RTs in valid (M=482.75, SEM=9.30) as compared to neutral (M=504.98, SEM=10.16) trials (t(34)=−5.97, p<.001), faster RTs in valid as compared to invalid (M=515.70, SEM=9.42) trials (t(34)=−6.88, p<.001) and a significant difference between neutral and invalid trials (t(34)=−2.37, p=.024) (figure 2B).

**Fig 2.**
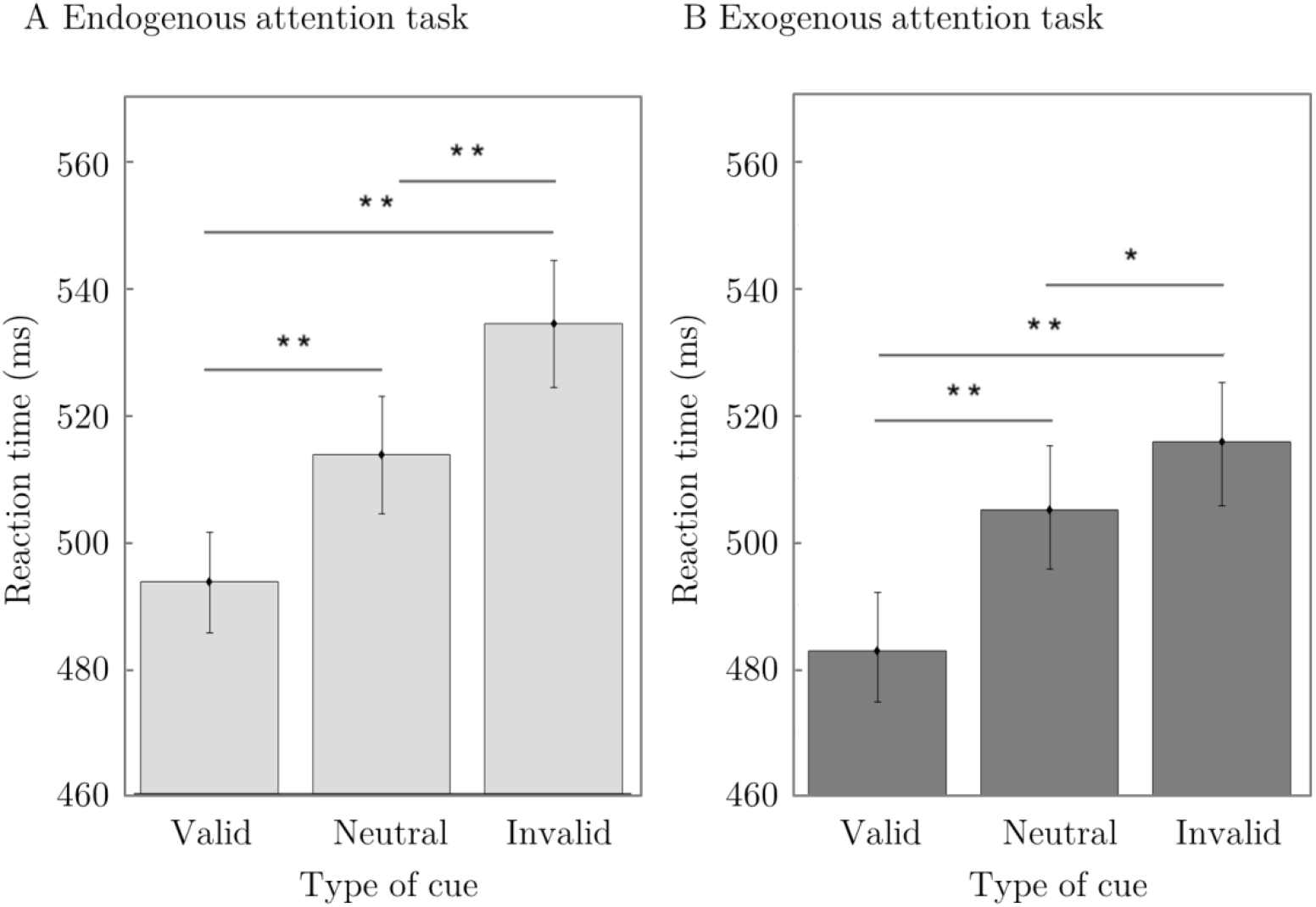
Cueing effect in the Endogenous and Exogenous attention task. A *RTs averaged over both hemifields per type of cue for the Endogenous attention task for the sham condition*. Participants’ reaction times were faster for valid cue trials as compared to neutral or invalid cue trials, and faster in neutral cue as compared to invalid cue trials, which indicates a successful cueing effect. B *RTs averaged over both hemifields per type of cue for the Exogenous attention task for the sham condition*. Validating the cueing effect, we found faster reaction times for valid cue trials as compared to neutral or invalid cue trials as well as faster reaction times in neutral cue as compared to invalid cue trials. One asterisk visualizes a significant difference with a p-value < 0.05 and a double asterisk indicates a significant difference with a p-value < 0.001.

### Visuospatial attention bias in the Endogenous attention task

We analyzed whether left parietal alpha tACS shifts attention to the left, by evaluating the effects of *Stimulation Condition* (tACS, sham) and *Type of Cue* (invalid, neutral, valid) on visuospatial attention bias (RT_left hemifield_ – RT_right hemifield_) in a Repeated Measures ANOVA. There was a main effect of *Stimulation condition* (F(1,33)=12.33, p=.001) with a greater leftward bias (M=9.29, SEM=6.30) in the 10Hz as compared to the sham condition (M=21.44, SEM=6.61) (figure 3A). The main effect of *Type of cue* (F(2,66)=.24, p=.790) and the interaction effect were not significant (F(2,66)=2.21, p=.118). The main effect of *Stimulation condition* confirms our hypothesis that left parietal tACS at alpha frequency induces a leftward bias in visuospatial attention relative to sham. Another way to present the same results is to subtract the sham (baseline) session data from the 10Hz tACS session data and subsequently compare the RTs between the two hemifields (figure 3B; statistically, this is identical to the main effect of stimulation condition). Here it can be seen that participants are faster for stimuli in the left (M=−12.58, SEM=10.94) as compared to the right hemifield (M=−.43, SEM=11.32).

**Fig 3.**
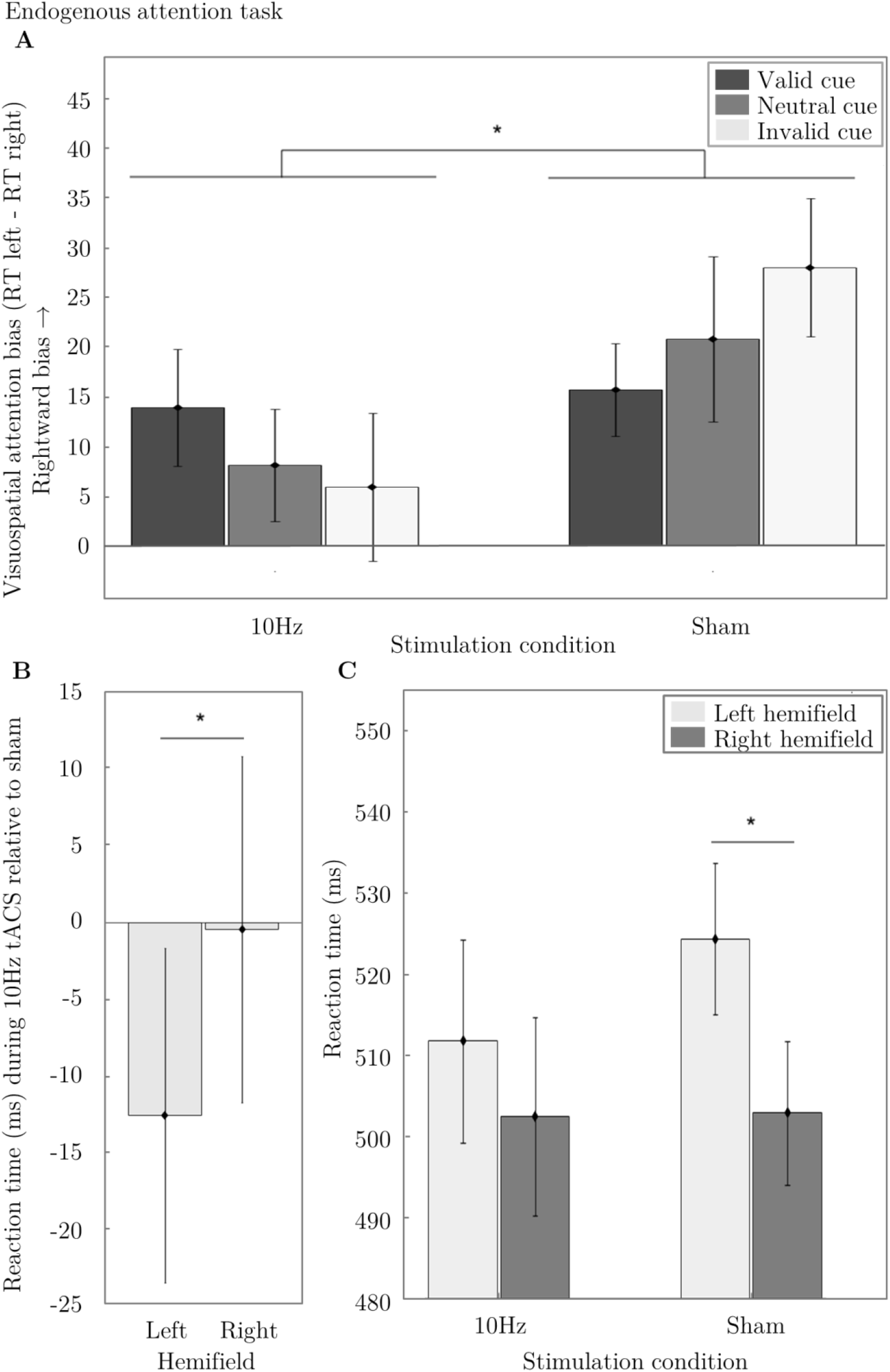
Results from the Endogenous attention task. A *Visuospatial attention bias in the Endogenous attention task during 10Hz and sham tACS for valid, neutral and invalid trials*. A positive value of visuospatial attention bias (RT_left hemifield_ - RT_right hemifield_) indicates a rightward bias whereas negative values indicate a leftward bias of visuospatial attention. There is a significantly greater leftward bias during 10Hz tACS as compared to sham. B *Reaction time per hemifield for the 10Hz stimulation condition relative to sham*. For 10Hz tACS relative to sham there is a difference in RTs for stimuli in the left (RT_left hemifield, 10Hz tACS_-RT_left hemifield, sham_) as compared to the right hemifield (RT_right hemifield, 10Hz tACS_ - RT_right hemifield, sham_).This means that we induced a leftward bias during 10Hz tACS relative to sham. Error bars visualize the standard error of the mean (SEM) across participants. C *RTs averaged over all trials per stimulation condition and hemifield*. For the sham stimulation condition, participants reacted faster in response to stimuli in the right as compared to the left hemifield. This effect is attenuated for the 10Hz stimulation condition. One asterisk visualizes a significant difference with a p-value < 0.05 and a double asterisk indicates a significant difference with a p-value < 0.001.

To test which hemifield drives the attentional bias effect, we analyzed the RTs averaged over all cues per *Hemifield* and S*timulation condition* in a Repeated Measures ANOVA. There was no main effect of *Stimulation condition* (F(1,33)=.35, p=.558), a main effect of *Hemifield* (F(1,33)=8.64, p=.006) and an interaction effect (F(1,33)=1 2.34, p=.001). In follow-up t-tests we found slower RTs for the left (M=524.41, SEM=9.37) as compared to the right hemifield (M=502.88, SEM=8.88) for the sham stimulation condition (t(33)=3.81, p=.002, *Bonferroni-corrected*). In contrast, the left hemifield (M=511.82, SEM=12.50) did not differ from the right hemifield (M=502.56, SEM=12.17) for the 10Hz stimulation condition (t(33)=1.72, p=.376, *Bonferroni-corrected*). The left hemifield of the 10Hz stimulation condition did not significantly differ from the left hemifield in the sham stimulation condition (t(33)=−1.15, p=1.0, *Bonferroni-corrected*). Likewise, the right hemifield of the 10Hz stimulation condition did not differ from the right hemifield of the sham stimulation condition (t(33)=−.04, p=1.0, *Bonferroni-corrected*) (figure 3C).

We thus found a greater leftward bias in the 10Hz stimulation condition as compared to sham in the Endogenous attention task, confirming our hypothesis. Follow-up analyses revealed that the RTs in neither the left nor right hemifield differed between the stimulation conditions. In the sham condition, there was a significant difference between hemifields with faster RTs to stimuli in the right as compared to the left hemifield. In the 10Hz condition, the two hemifields did not differ from each other. However, when subtracting the data of the sham from the 10Hz tACS condition, we found a leftward bias with faster RTs for the left as compared to the right hemifield.

### Visuospatial attention bias in the Exogenous attention task

To test whether tACS at 10Hz induced a leftward bias in visuospatial attention relative to sham we ran a Repeated Measures ANOVA with visuospatial attention bias as dependent variable and *Stimulation condition* (10Hz and sham) and *Type of cue* (Valid, Neutral, and Invalid) as factors. There was a main effect of *Type of cue* (F(2,68)=6.61, p=.002) but neither a main effect of *Stimulation condition* (F(1,34)=2.25, p=.143) nor an interaction effect (F(2,68)=.50, p=.610) (figure 4A). This means that tACS at 10Hz did not induce a significant leftward bias relative to sham.

**Fig 4.**
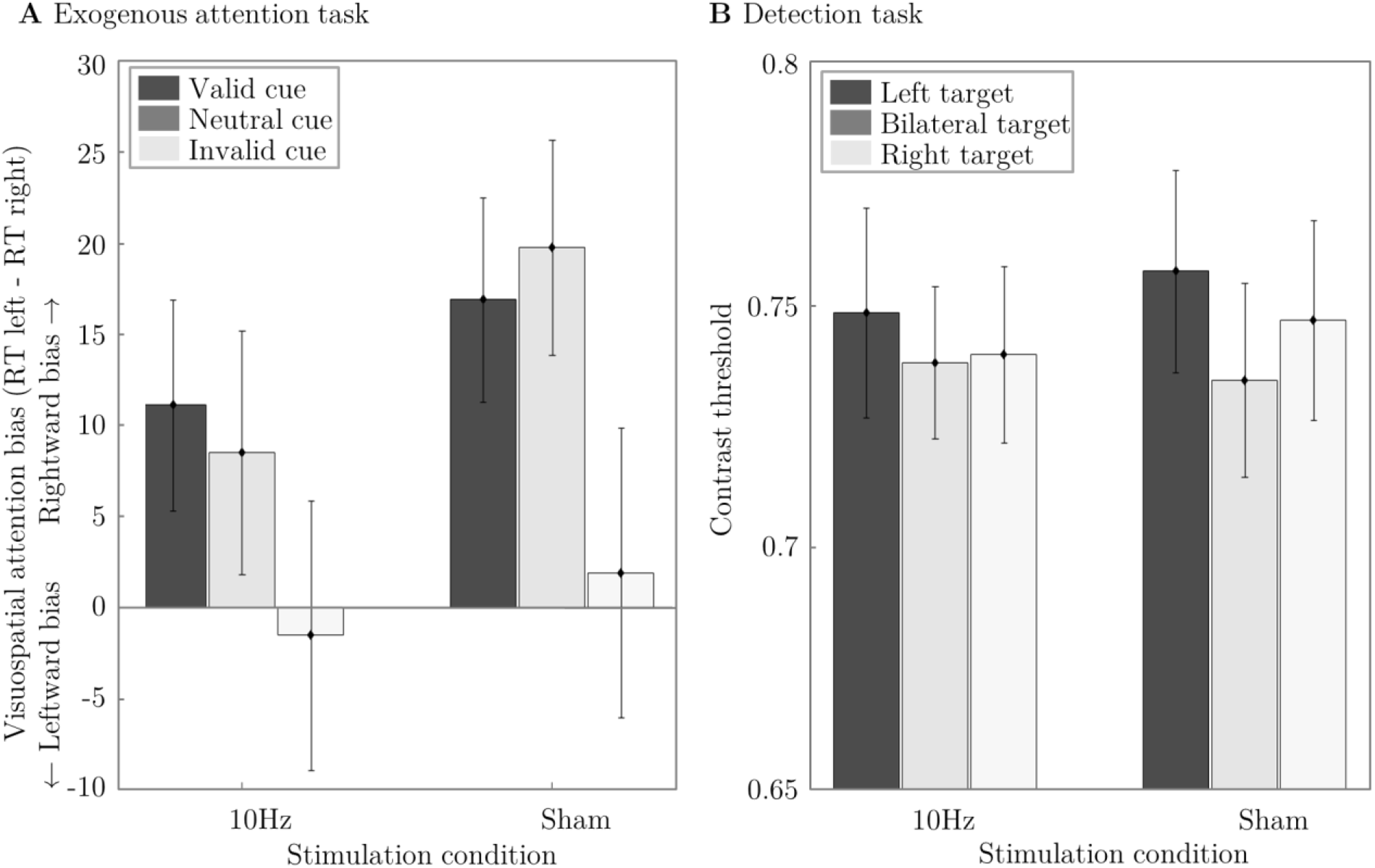
Results from the Exogenous attention and Detection tasks. A *Visuospatial attention bias in the Exogenous attention task during 10Hz and sham tACS for valid (dark grey), neutral (grey) and invalid (light gray) trials*. There was no main effect of Stimulation Condition or interaction effect but only an effect of Cue Type. B *Contrast thresholds in the Detection task during 10Hz and sham tACS for left (dark grey), bilateral (grey) and right (light grey) targets*. There were no significant main or interaction effects. Error bars visualize the standard error of the mean (SEM) across participants.

Similar to the analysis of the Endogenous attention task, we analyzed the RTs averaged over all cues per *Hemifield* and *Stimulation condition* in a Repeated Measures ANOVA. There were no significant effects (*Stimulation condition:* (F(1,34)=.03, p=.857), *Hemifield:* (F(1,34)=3.34, p=.077), *Stimulation Condition * Hemifield:* (F(1,34)=2.25, p=.143). Hence, 10Hz tACS did not significantly affect visuospatial attention performance in the Exogenous attention task

### Visuospatial attention bias in the Detection task

A repeated Measures ANOVA with *Stimulation condition* (10Hz or sham) and *Hemifield* (left, right or both hemifields) as factors and contrast thresholds as dependent variable did not reveal a main effect of *Stimulation condition* (F(1,35)=.06, p=.813), *Hemifield* (F(2,70)=.94, p=.397) or an interaction effect (F(2,70)=.284, p=.754) (figure 4B). To test whether tACS at 10Hz induced extinction-like effects we analyzed the errors for the bilateral trials, i.e. whether participants only perceived the left or the right stimulus when actually a bilateral stimulus was shown. *Stimulation Condition* and *Indicated location of the stimulus* (left or right) was added as factors in a Repeated Measures ANOVA and number of correct trials as dependent variable. There was no main effect of *Stimulation Condition* (F(1,35)=.53, p=.471) or *Indicated location of the stimulus* (F(1,35)=.16, p=.695) and no interaction effect (F(1,35)=.64, p=.429). This suggests that performance in the detection task was not affected by tACS at 10Hz.

## Discussion

Previous EEG studies have shown an association between bias in endogenous attention and lateralization of occipitoparietal alpha power, showing greater power in the ipsilateral relative to the contralateral side of attention [4–7], Here, we tested whether this association is robust enough to permit manipulations of the spatial distribution of attention by experimentally inducing hemispheric increases in oscillatory alpha power using NIBS. We therefore stimulated the left parietal cortex with high-density tACS either at 10Hz or sham while assessing the bias in visuospatial attention with an Endogenous and Exogenous attention task, as well as a visuospatial detection task. The present report is (among) the first to show that visuospatial attention can be influenced by tACS at alpha frequency. In the Endogenous attention task, a robust leftward bias was induced during 10Hz tACS as compared to sham. Interestingly, no significant stimulation effects were found in the detection task and Exogenous attention task, indicating a task-specificity of the left parietal tACS intervention.

### Task specific tACS effects

A tACS-induced leftward bias was found in the endogenous but not in the Exogenous attention task or detection task. The absence of stimulation effects for the exogenous cueing task in our experiment might speak against an association between exogenous attention and parietal alpha oscillations. The commonly reported lateralization of alpha power is observed after the presentation of an endogenous cue but before a target stimulus is shown [4–7], This anticipatory change in hemispheric alpha power prior to target onset speaks in favour of an endogenous rather than an exogenous attention process. Alternatively, it is possible that the exogenous cueing effects simply outweighed the tACS effect. It has previously been shown that voluntary orienting can be interrupted by salient lateralized cues [36–38], Accordingly, the tACS induced endogenous attention bias in our experiment may have been overruled by the exogenous cues.

Analyzing the contrast thresholds of the detection task, there was also no evidence for a leftward bias during 10Hz tACS as compared to sham. In contrast to the orienting tasks, which involved an orientation discrimination task and attentional manipulations with cues, the detection task simply measured low-level perceptual sensitivity without attentional manipulation. Our findings are in line with a recent transcranial current stimulation (tDCS) reporting no effect of parietal tCS on contrast thresholds [3]. It could be argued that left parietal tACS did not affect lower-level visual processing (e.g. target detection performance) but rather higher-level attentional processes. However, it has previously been shown that within and between subject target detection performance is associated with pre-stimulus occipitoparietal alpha power, which stands in contrast to our findings [39,40]. In those experiments, the stimulus was presented in the center of the screen and the change in alpha power was measured at medial electrode sites. Here, we used lateralized target stimuli and stimulated the left parietal cortex, which might explain the absence of stimulation effects in the detection task for our experiment. Alternatively, it could be argued that our detection task was not sensitive enough to measure the visuospatial attention bias induced by tACS at alpha frequency.

### Stimulation site and ring electrodes

Through left parietal tACS at 10Hz, we successfully shifted visuospatial attention to the left hemifield. In a similar attempt, Veniero and colleagues [31] conducted two consecutive studies in which they targeted the right parietal cortex with tACS at alpha frequency in order to induce a rightward bias. In experiment 1, they found the expected rightward bias for 10Hz tACS as compared to sham in a line bisection (landmark) task, but this finding could not be replicated in a second experiment. Likewise, Hopfinger, Parsons and Froehlich [41] administered right parietal tACS at alpha frequency but report no visuospatial attention bias as compared to sham in an Endogenous and Exogenous attention task.

These results are surprising considering the rather established association of parietal alpha power lateralization with visuospatial attention. What distinguishes our experiment from the above mentioned studies is that we used ring instead of disc electrodes. Compared to standard, rectangular, electrode configurations, ring electrodes enable a higher spatial focality [34], making it possible to limit stimulation to the left parietal cortex. Another difference to the above mentioned studies lies in the stimulation site. We stimulated the left instead of the right parietal cortex. A recent fMRI experiment including an Endogenous as well as an Exogenous attention task showed that task-related activity is greater in the left as compared to the right frontoparietal attention network [42]. Here, the left hemisphere seemed to be especially involved in reorienting, showing greater activation for invalid as compared to valid trials. Moreover, the change in functional connectivity during an endogenous attention task as compared to rest has shown to be more pronounced in the left as compared to the right hemisphere [43]. At rest, functional connectivity in the frontoparietal network was tonically higher in the right as compared to the left hemisphere. However, the left hemisphere was more specifically recruited during high attentional demands thereby balancing out the right hemispheric asymmetry. This might explain why left parietal tACS in our experiment induced a leftward bias in visuospatial attention whereas right parietal tACS has previously led to inconsistent results.

### Clinical relevance and suggestions for future research

The possibility of modulating alpha oscillations through tACS at alpha frequency is not only relevant in the framework of fundamental research but might also have implications for the treatment of hemineglect patients. Common rehabilitation treatments for neglect patients focus on the contralesional enhancement of attention [44] through e.g. prism adaptation [45] or vestibular stimulation [46,47]. Recently, transcranial magnetic stimulation approaches have been introduced for non-invasively disrupting the unaffected hemisphere and thereby alleviating neglect symptoms [48,49]. In the present report, we showed that it is possible to induce a visuospatial attention bias in healthy participants through unilateral parietal tACS at alpha frequency. In order to verify that the reported stimulation effects are frequency specific and limited to the alpha range, future research should also include a control stimulation frequency condition. Another focus should lie on the individualization of stimulation protocols using individual stimulation frequencies. According to general theories of en-trainment and network models, this should boost the effects [50,51] and might become especially important when applying tACS in a heterogeneous group of patients. It remains to be seen if tACS at alpha frequency can also be used to treat patients with attentional deficits.

Hemineglect patient commonly suffer from a pathological rightward bias [8,9]. This rightward bias could be counteracted with left parietal tACS at alpha frequency, as demonstrated here with healthy participants. TACS has been proposed to induce neuroplastic changes under the stimulation site [52–54], This might make it a potential easy-to-apply, portable and affordable treatment for hemineglect patients with longterm benefits [52].

## Acknowledgements

We would like to thank Jeannette Boschma for helping with the data collection. This work was supported by the Netherlands Organization for Scientific Research (NWO, Veni to T.G. 451-13-024; Vici to A.S. 453-15-008).

